# Endothelial Colony-Forming Cell Transcriptomic Profiling in CT-defined Coronary Artery Disease from the BioHEART-CT Study Implicate CCBE1 in Mitochondrial Dysfunction-associated Atherosclerosis

**DOI:** 10.1101/2025.08.18.670989

**Authors:** W. Eugene Lee, Sarah L. Cook, Gareth S.D. Purvis, Albert Henry, Kirsten Schnitzler, Elijah Genetzakis, Marie Besnier, Madhuvanthi Chandrakanthan, Andy Tran, Dario Strbenac, Carmen Mifsud, Katharine Bate, Stephen T. Vernon, Tung V. Nguyen, Michael P. Gray, Stuart M. Grieve, Jean Yang, Daniel MacArthur, Joseph E. Powell, Keith M. Channon, Gillian Douglas, Gemma A. Figtree

## Abstract

**Background:** Endothelial dysfunction is an early contributor to atherosclerosis. This study combined CT imaging of coronary artery disease (CAD) and patient-dervied endothelial colony-forming cells (ECFCs) transcriptional profiling to investigate potential mechanisms underlying endothelial dysfunction in atherosclerosis.

**Methods:** Twenty-six individuals with CT-defined CAD and eighteen non-CAD controls were included in the Discovery Cohort for bulk RNA sequencing and transcriptomic analysis of ECFCs. Differential gene expression analysis was performed, and candidate genes were selected based on logFC and p-value. Candidate genes were carried forward for gene expression validation using quantitative real-time PCR (qRT-PCR) in a Validation Cohort. Mitochondrial reactive oxygen species (mROS) production and mitochondrial mass were assessed using multi-colour flow cytometry. Functional validation of the top candidate was conducted in using human umbilical vein endothelial cells (HUVECs) using loss-of-function genetic approach. Expression Quantitative Trait Loci (eQTL)-association analysis was conducted using genotype data from the BioHEART-CT cohort.

**Results:** Pairwise analysis identified six differentially expressed protein-coding genes in CAD ECFCs: *CCBE1 (Collagen and Calcium Binding EGF Domain-Containing 1), SPINT2, CRISPLD1, PIEZO2, EPB41L3*, and *AC005943.1*. qRT-PCR in the Validation Cohort confirmed significantly higher *CCBE1* expression in CAD patients. Individuals with relative *CCBE1* fold change expression>10 had a 2.8-fold increase in the log-odds ratio of CT-defined CAD. CAD ECFCs displayed elevated mROS and mitochondrial mass. *CCBE1* knockdown in HUVECs reduced mROS and mitochondrial mass without affecting proliferation or permeability, but shifted cells into a metabolically elevated state, marked by increased ATP production, respiration and glycolysis. *CCBE1* cis-eQTLs were associated with increased odds of CAD in the BioHEART-CT cohort.

**Conclusions:** *CCBE1* expression in ECFCs was higher in patients with CT-defined CAD versus non-CAD. Quantitative assessment of transcript levels supported a causal relationship between greater *CCBE1* expression and CAD burden and risk, and functional experiments on *CCBE1* knockdown demonstrated improved mitochondrial function in human endothelial cells.

**GRAPHICAL ABSTRACT:** 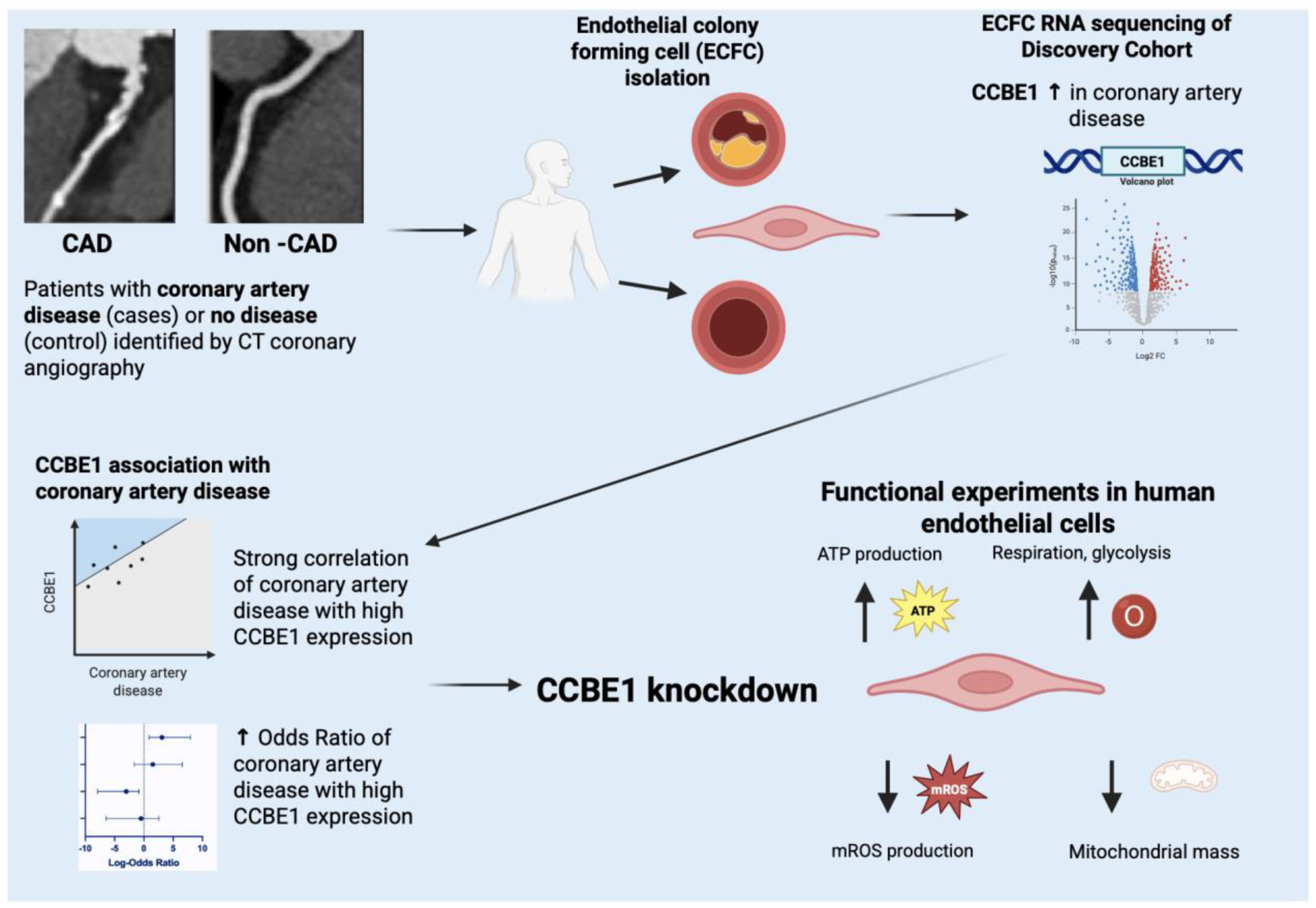

**Novelty and Significance:** *What is Known?:*

- The endothelium plays a critical role in vascular health and susceptibility to atherosclerosis.
- Mitochondrial dysfunction has been implicated in atherosclerosis, but its role and mechanism in individual susceptibility to CAD in humans is not known.

*What New Information Does This Article Contribute?:*

- Novel approach integrating CT imaging with ECFC functional data to link vascular structure with endothelial biology *ex vivo*.
- Patients with CT-defined CAD had 3.6-fold higher *CCBE1* expression compared to non-CAD within the Validation Cohort.
- *Cis*-eQTL-association analysis revealed increased odds of CAD.
- *CCBE1* knockdown improved mitochondrial function in human endothelial cells.
- Together, these 4 lines of evidence point to a novel and causal role for *CCBE1* in human CAD.

## Introduction

Over the past five decades, our understanding of coronary artery disease (CAD) has been greatly influenced by the identification and targeting of well-established risk factors such as diabetes, smoking, hypertension, and hypercholesterolemia^1, 2^. While these efforts have undoubtedly contributed to advancements in treatment and prevention strategies, a significant proportion of first-time heart attack patients (up to 27%) have developed their underlying CAD despite having none of these standard modifiable risk factors (SMuRFs)^3, 4^. This phenomenon underscores a critical gap in our current approach, as it suggests that there are underlying mechanisms driving the progression of heart disease beyond those currently recognised.

We have previously reported that patients at heightened susceptibility to CAD and ST-elevation MI against a “zero SMuRF” background have an almost 50% higher 30-day mortality compared to those with conventional risk factors^5–9^. In addition to underscoring the critical need for early diagnosis and preventive strategies in this population, these findings prompt important mechanistic questions regarding the biological drivers of their heightened susceptibility and increased incidence of major adverse cardiac events (MACE). Heightened or dysregulated oxidative signalling and/or inflammation are strong candidates for this^10^. Unbiased approaches to discovery dysregulated molecular pathways in these individuals may provide novel insights into CAD mechanisms that may benefit the broader population in regard to diagnostic and therapeutic tools^9^.

Whilst numerous teams have established biobanks for discovery of new mechanisms and markers of CAD and associated clinical events, most have focused on the more accessible and pragmatically feasible biosamples such as DNA, serum and plasma. Endothelial cells (ECs) play a pivotal role in maintaining vascular homeostasis, and dysfunction in these cells has been implicated in the initiation and progression of atherosclerosis, the hallmark of CAD^11–16^. To delve deeper into the mechanisms underlying endothelial dysfunction in CAD, we have developed a protocol for culturing endothelial colony forming cells (ECFCs) from the blood of patients with and without CAD defined by CT-coronary angiography (CTCA)^17, 18^. We have established a subcohort of the BioHEART-CT study (Australia New Zealand Clinical Trials Registry ANZTR12618001322224, Camperdown, Australia) comprising over 600 ECFC lines from patients with CTCA, and reported a persistant phenotype reflecting CAD^17^. Previously, we found a strong correlation between mitochondrial reactive oxygen species (mROS) production and CAD severity in these ECFCs, by quantitative measures of coronary artery calcium score (CACS) or Gensini score^18^. This has led to efforts by our team to translate measures of ECFC mROS production as a potential model of human CAD susceptibility using high-throughput, high-content mROS screening systems^19^.

Building on our prior characterisation of ECFCs from CAD patients, we now turn to an unbiased transcriptomic approach to further investigate molecular mechanisms underlying CAD. By profiling gene expression in ECFCs from patients with and without CAD-as defined by CTCA, within the BioHEART-CT cohort, we aim to identify dysregulated genes and pathways that may drive disease susceptibility. Our objective is to identify novel molecular signatures specific to CAD-affected ECs and gain insights into cellular functions that could be therapeutically targeted to slow or prevent disease progression. Through this approach, we identified CCBE1 (Collagen and Calcium Binding EGF Domain-Containing 1) as a novel candidate gene enriched in CAD ECFCs, with potential roles in modulating mitochondrial function.

## Methods

### Data Availability

The data that support the findings of this study are available from the corresponding authors upon reasonable request. All methods are described in detail in the Supplemental Material.

## Results

### Characteristics of the BioHEART-CT Cohort

The demographics of the BioHEART-CT patients with CT-defined coronary atherosclerosis burden and matching ECFCs cultured from individual patient peripheral blood are presented in Table 1. Males represented a greater proportion in the CAD cohort compared to the non-CAD cohort (65.4% versus 38.9% respectively). Notably, 26.9% of the CAD patients in the Discovery Cohort, and 21.1% of the CAD patients in the Validation cohort had developed coronary atherosclerosis despite no SMuRFs. This is similar to the proportion of ST-elevation MI patients at our centre with confirmed atherosclerotic culprit lesion despite having no SMuRFs^8^.

**Table 1.**
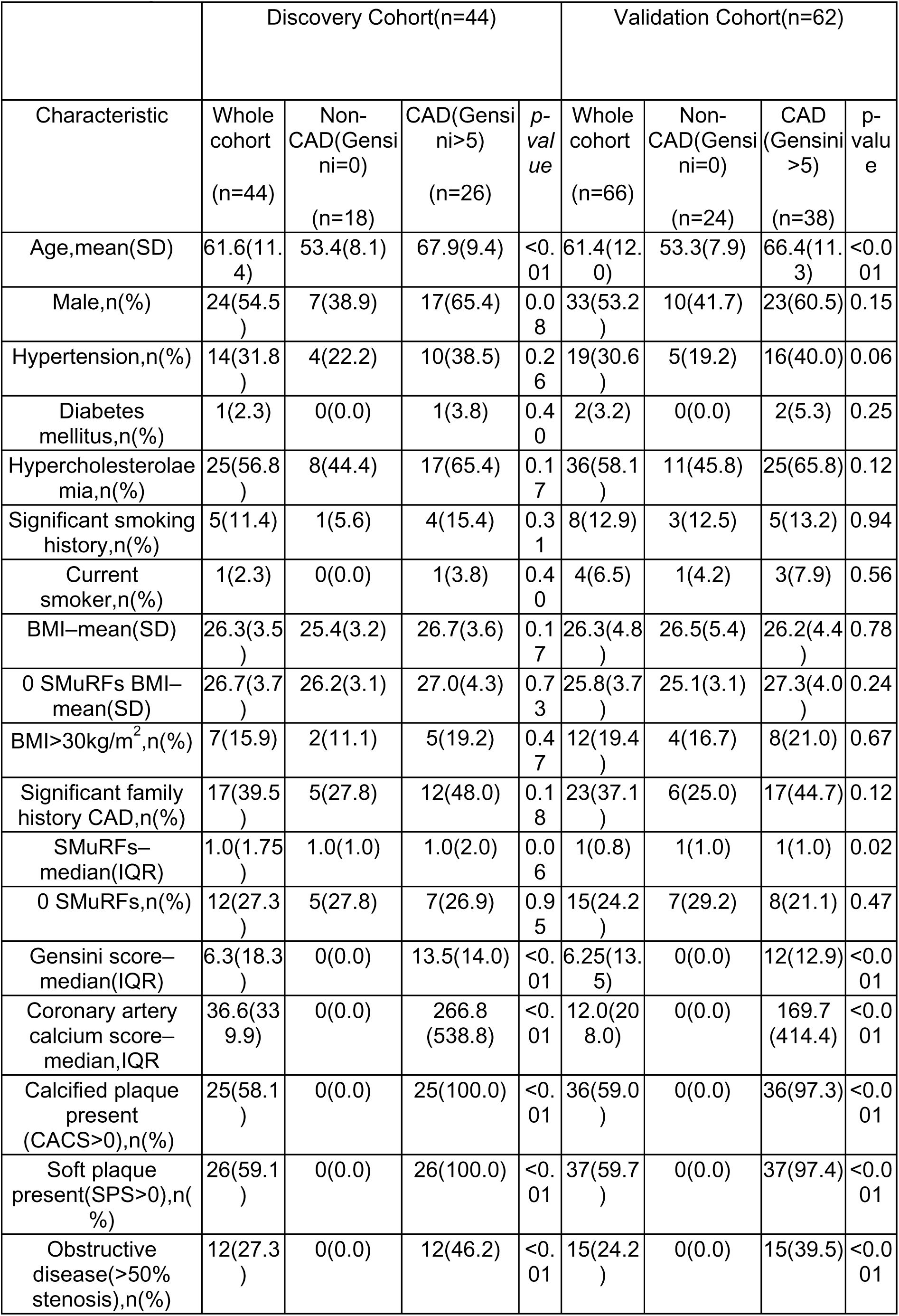
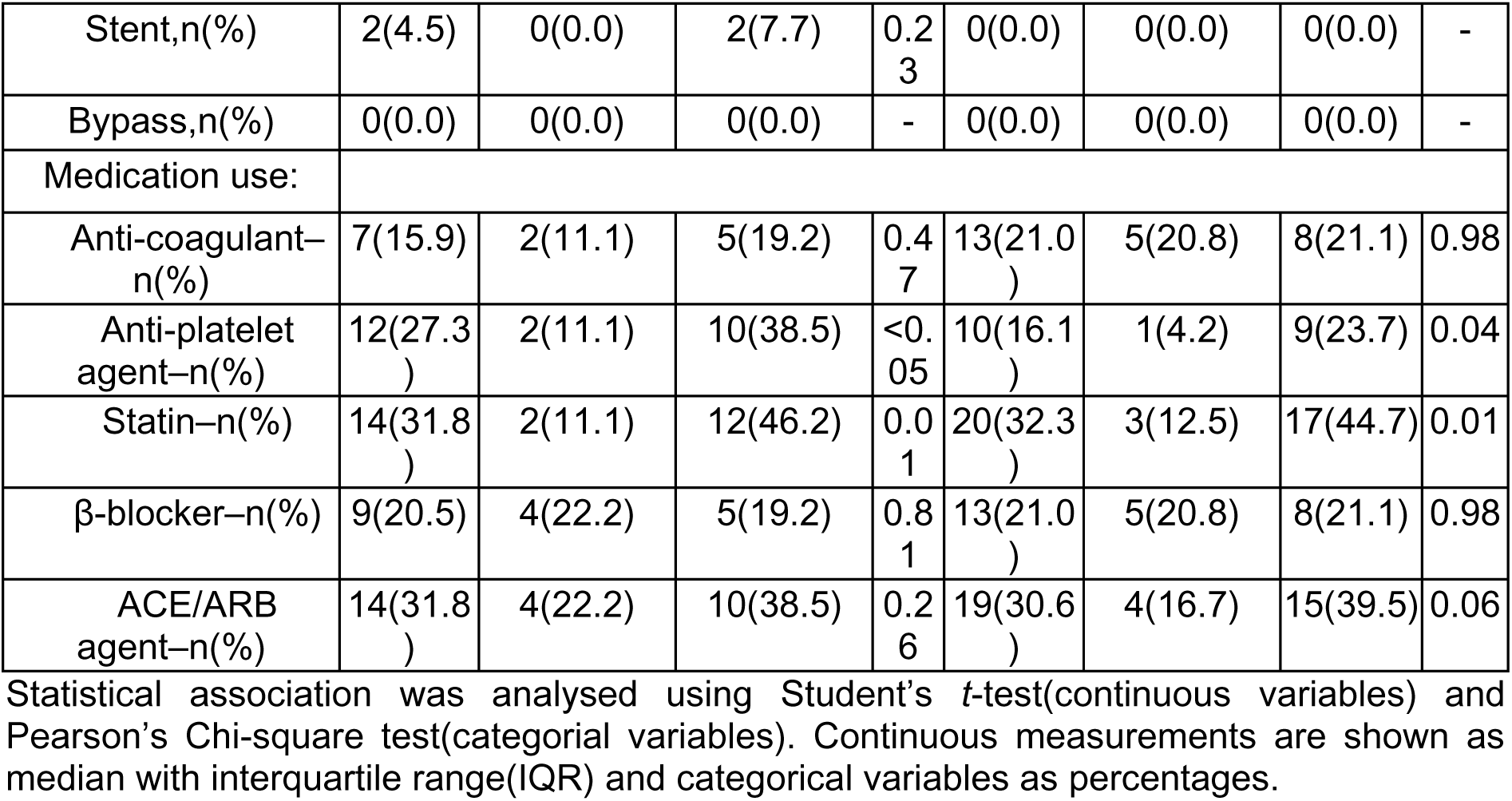
Demographical and clinical characteristics of the BioHEART-CT cohorts.

### Transcriptomic profiling of ECFCs identifies CAD-linked DEGs and pathways involved in oxidative stress

We analysed the transcriptional profile in ECFCs from patients with or without CAD in the discovery cohort (Figure 1A). 12, 725 genes were identified and pairwise analyses yielded 7 differentially expression genes (DEGs) with absolute log_2_FC≥1 and unadjusted P<5.0×10^−2^ which represents 0.06% of the total transcriptome (Figure 1B). Specifically, the expression of *SPINT2 (Kunitz-type protease inhibitor 2), CRISPLD1 (Cysteine rich secretory protein LCCL domain containing 1), CCBE1 (Collagen and calcium-binding EGF domain-containing protein 1), PIEZO2 (Piezo-type mechanosensitive ion channel component 2), EPB41L3 (erythrocyte membrane protein band 4.1 like 3)* and *AC005943.1* were increased whereas *RFLNA* (*Refilin A*) expression was decreased (Figure 1B;Supp. Table S1).

**Figure 1.**
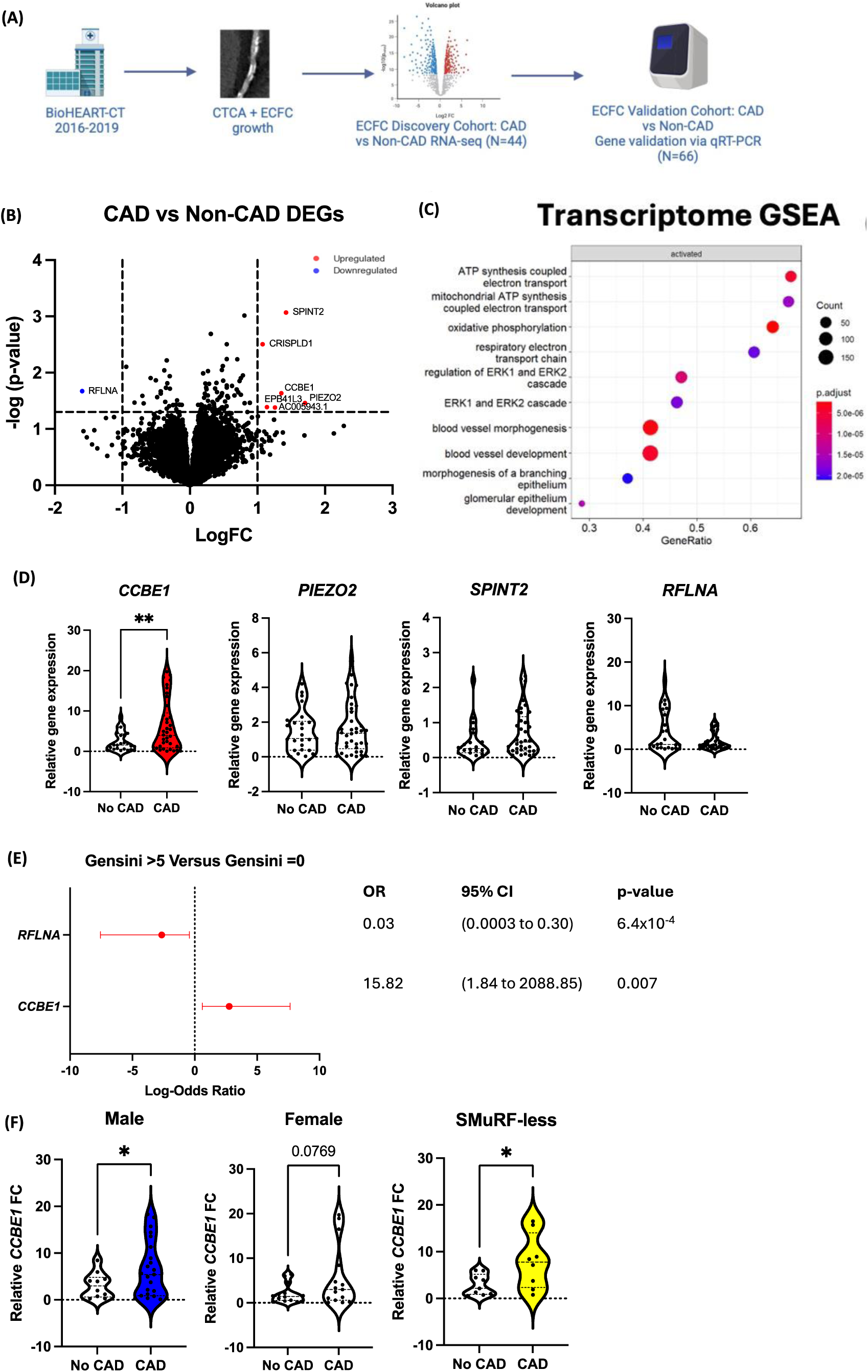
Patient-derived ECFC Transcriptome reveal mitochondrial-associated transcriptional profile, with CCBE1 being differentially expressed. The presence of CAD was determined using Gensini>5 is CAD and Gensini=0 is Non-CAD.(A)Schematic diagram of the workflow for BioHEART-CT ECFC transcriptomic analysis.(B) Volcano plot of DEGs from bulk RNA-seq data of CAD and non-CAD ECFCs.(D)Enrichment analysis of DEGs identified by bulk RNA-seq show GO terms representative of mitochondrial pathways from CAD-ECFCs compared to non-CAD-ECFCs;N=44(Non-CAD=18, CAD=26).(D)Validation of DEGs in patient ECFCs by qRT-PCR. Results are expressed as relative FC from reference sample between CAD and no CAD.Results are presented as violin plots.(E)LogOR for the association between *CCBE1* (FC>10) and CAD from a univariate model. Error bars represent 95% CI. Corresponding OR with 95% CI are presented in adjacent table. (F)Subcohort differences in *CCBE1* expression between male, female and SMuRF-less subcohorts. Results are presented as violin plots. Statistical analyses were performed using independent samples t-test or Firth’s bias-reduced penalized likelihood method for logistic regression test.*P<5×10^−2^, **P<1.0×10^−2^.DEG, differentially expressed genes;FC, fold change;LogOR, log-odds ratio;CI, confidence intervals;SMuRF, standard modifiable risk factors.

Gene Set Enrichment Analysis (GSEA) identified 696 GO terms after false discovery rate (FDR) correction, with the top 10 terms (FDR<1.0×10^−2^) based on gene ratio shown in Figure 1C and Supp. Table S2, revealing a significant association between CAD and enriched pathways involved in ATP synthesis and the electron transport chain (ETC). Additionally, 26 KEGG pathways were enriched, with the top 10 pathways highlighting key biological processes such as oxidative phosphorylation, which drives ATP production, and ROS generation, a key contributor to oxidative stress (Supp. Figure 1A;Supp. Table S3).

### CAD ECFCs have upregulated *CCBE1* expression

To validate our findings, we performed qRT-PCR on the protein-coding DEGs present in the top 5 CAD-associated DEG list (*CCBE1, PIEZO2, RFLNA, SPINT2*;Supp. Table S1). Among these, *CCBE1* was significantly upregulated in CAD patients compared to non-CAD patients (mean FC difference:3.66-fold, P<5.0×10^−3^;Figure 1D). We observed a bimodal distribution within the CAD group; with a subset of CAD patients expressing high *CCBE1* (relative FC>10) and the other CAD subgroup expressing low *CCBE1* (relative FC<10). Of interest, not a single patient from the non-CAD group had high *CCBE1*. No differences were found between the patients’ SMuRFs. Using this visible cut-off, high *CCBE1* was associated with increased odds of CAD (odds ratio(OR):15.82;95% CI:1.84-2088.85;P=7.0×10^−3^;Figure 1E). Sex diaggregated analysis was performed where male CAD patients were found to have significantly upregulated *CCBE1* expression (mean FC difference:3.36-fold, n=33;P=3.4×10^−2^) compared to their non-CAD counterparts (Figure 1F). Whilst a similar trend was observed in the female sub-cohort, this was not significant, limited by the number of participants (n=29;P=8.0×10^−2^). Within the SMuRF-less CAD subcohort, a significant increase (mean FC:4.99, P=3.0×10^−2^) was observed compared to their non-CAD counterparts. CCBE1 protein expression was also explored however no differences were detected (Supp. Figure 1B-C).

Similarly, *RFLNA* was observed to have a bimodal distribution in the non-CAD group despite no difference between CAD and non-CAD patients (Figure 1D). A subset had high *RFLNA* (relative FC>6.35), while others had low *RFLNA* (relative FC<6.35). Using this threshold, high *RFLNA* was associated with a reduced odds for CAD (OR:0.03;95% CI:0.0003-0.30;P=6.4×10⁻⁴;Figure 1E).

### *CCBE1* Expression in Patient ECFCs have Strong Associations for Absolute and Relative CAD burden

From the results above, we aimed to investigate the relationship between *CCBE1* expression and the most common clinical measure of coronary artery calcium-the Agatston score^20^. We assessed *CCBE1* in relation to clinically actionable CAD metrics, both through absolute burden (CACS>100 Agatston unit(AU)) and relative risk burden (CACS percentile>75%)^21, 22^. *CCBE1* levels were significantly associated with absolute CACS burden (median FC difference:2.34, P<5.0×10^−2^;Figure 2A) and with CACS percentile (median FC difference:3.32, P<5.0× 10^−3^;adjusted using the MESA formula for age and gender;Figure 2B). A significant positive correlation was found between *CCBE1* FC and Log(CACS+1) (R=0.35, P=3.0×10^−3^;Figure 2C), and with CACS percentile (R=0.28, P=1.8×10^− 3^;Figure 2D). Sex disaggregated analyses showed no interaction between sex and *CCBE1* in regard to CACS association (Supp. Table S4-5).

**Figure 2.**
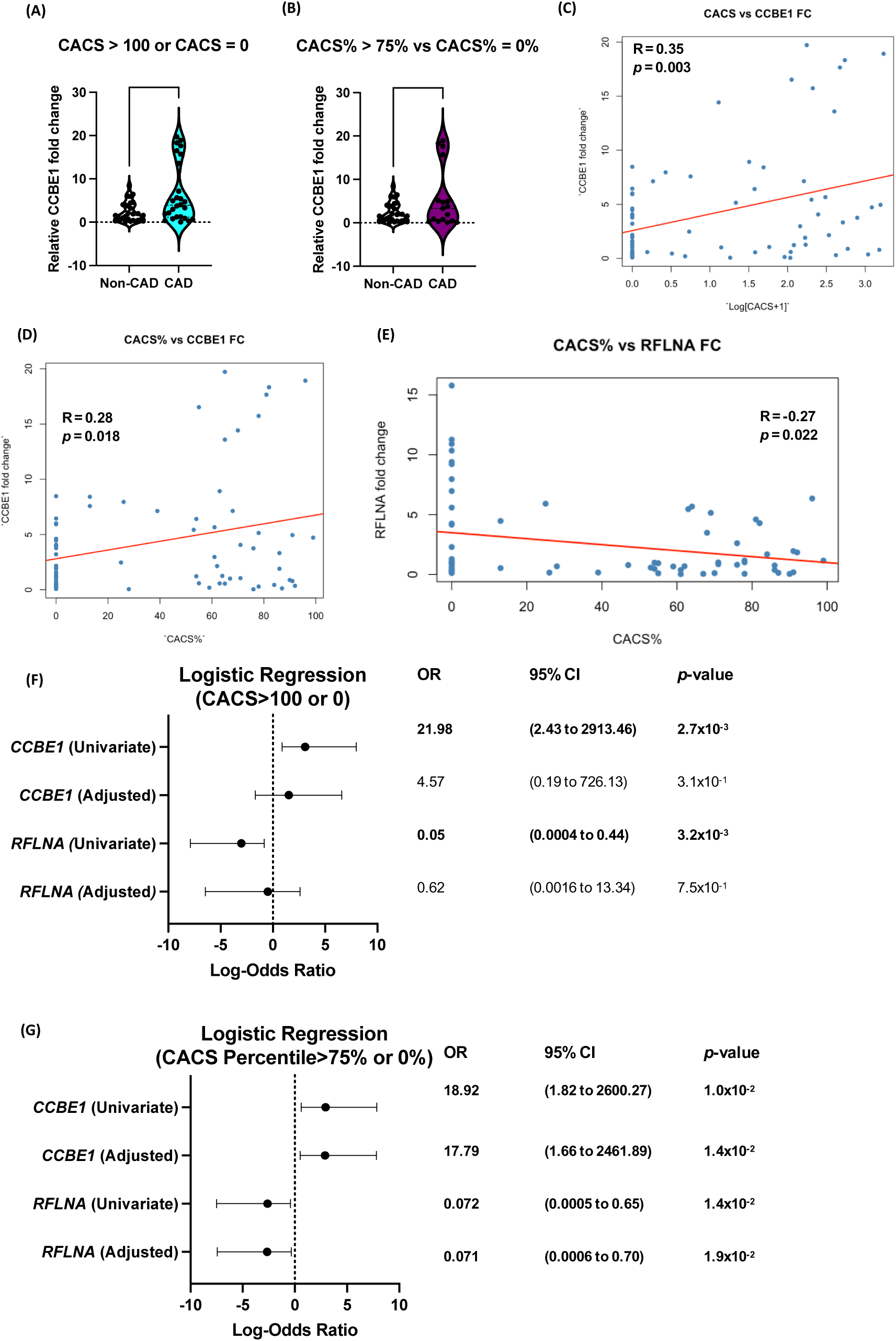
High *CCBE1* Expression (FC>10) Strongly Predicts clinically actionable CAD. (A)Comparison of *CCBE1* FC levels with absolute CACS>100AU vs. CACS=0AU and (B)CACS percentile>75% vs. CACS percentile=0%, showing a significant difference in both CAD and non-CAD groups. Results are expressed as violin plots. Simple linear regressions of *CCBE1* FC against (C)Log[CACS+1] and (D)CACS%.(E)Simple linear regression of *CCBE1* fold change against Log[CACS+1] in female patients. LogOR of (F)CACS>100 AU vs. CACS=0 AU and (G)CACS percentile>75% vs. CACS percentile=0% with *CCBE1* FC>10. Logistic regression was adjusted for age, sex and SMuRF covariates for CACS whilst only SMuRF covariates were adjusted for CACS percentile. Corresponding OR with 95% CI are presented in adjacent table. Statistical analyses were performed using independent samples test or Firth’s bias-reduced penalized likelihood method for logistic regression test.*P<5×10^−2^, **P<1.0×10^−2^.AU, Agatston Unit;CACS, coronary artery calcium scoring;FC, fold change;LogOR, log-odds ratio;SMuRF, standard modifiable risk factors.

No significant differences were observed between *RFLNA* relative FC and CACS>100AU or >75% percentile (Supp. Figure 2A-B). While *RFLNA* showed no association with absolute CACS burden (Supp. Table S6), it was negatively correlated with CACS percentile (R=-0.27, P=2.2×10⁻²;Figure 2E). Sex-disaggregated analyses demonstrated no interaction between sex and *RFLNA* levels for CACS association (Supp. Tables S6-7).

Similarly to previously, we used the observed cut-point for high *CCBE1* to examine the association or potential prognostic ability of *CCBE1* above these levels. Individuals with high *CCBE1* had increased odds of absolute CACS>100AU (OR:21.98;95% CI:2.43-2913.46, P=2.7×10^−3^;Figure 2F). High *CCBE1* also increased odds of CACS percentile>75% (OR:18.92;95% CI:1.82-2600.27;P=1.0×10^−2^) which remained after SMuRF covariates (diabetes, hypertension, hypercholesterolemia, smoking status)-adjustment (OR:17.79;95% CI:1.66-2461.89;P=1.4×10^−^2;Figure 2G). Sex-specific and SMuRFless analyses are presented in Supp. Fig 2C-D.

Patients with high *RFLNA* had significantly lower odds of absolute CACS>100AU (OR:0.05;95% CI:0.0004-0.44, P=3.2×10⁻³;Figure 2F). Similarly, high *RFLNA* was associated with decreased odds of CACS percentile>75% (OR:0.07;95% CI:0.0005-0.65;P=1.4×10⁻²), which persisted after adjusting for SMuRF covariates (OR:0.07;95%CI:0.0006-0.70;P=1.9×10^−2^;Figure 2G). Sex-specific and SMuRFless cohort analyses are provided (Supp. Figures 2E–F). Given our interest in CAD-associated gene mechanisms, the remainder of this manuscript focuses on the mechanistic studies of *CCBE1*.

### CAD ECFCs have increased mitochondrial mass and mROS production

Building on the transcriptome-wide GSEA and KEGG analyses that highlighted mitochondrial and oxidative stress pathways, and given the strong associations between mROS generation and CAD severity - as defined by Gensini score and CACS^18, 19^–we investigated mROS production in parallel with mitochondrial mass, a known modulator of ROS levels^23^. This integrative approach enabled a more nuanced characterisation of ECFCs (Figure 3A-B). Here, a higher proportion of CAD ECFCs (Gensini>5, CACS>100AU or >75% percentile) exhibited high mROS-generation (MitoSOX Red^high^) and high mitochondrial-mass (MitoTracker Green^high^;Figure 3C). Compared to Gensini=0, the Gensini>5 group had 28.79% more high mROS-generating ECFCs (P=1.0×10⁻²) and 29.20% increased mitochondrial-mass (P=8.6×10⁻³). Similarly, CACS>100 AU patients had 28.00% more high mROS-generation ECFCs (P=2.8×10⁻²) and 29.96% higher mitochondrial-mass (P=6.4×10⁻³) than CACS=0 patients. Patient ECFCs from CACS percentile>75% had 29.08% more high mROS-producing cells (P=1.5×10^−2^) and 34.21% more high mitochondrial-mass ECFCs than those with 0% percentile (P=3.2×10^−3^).

**Figure 3.**
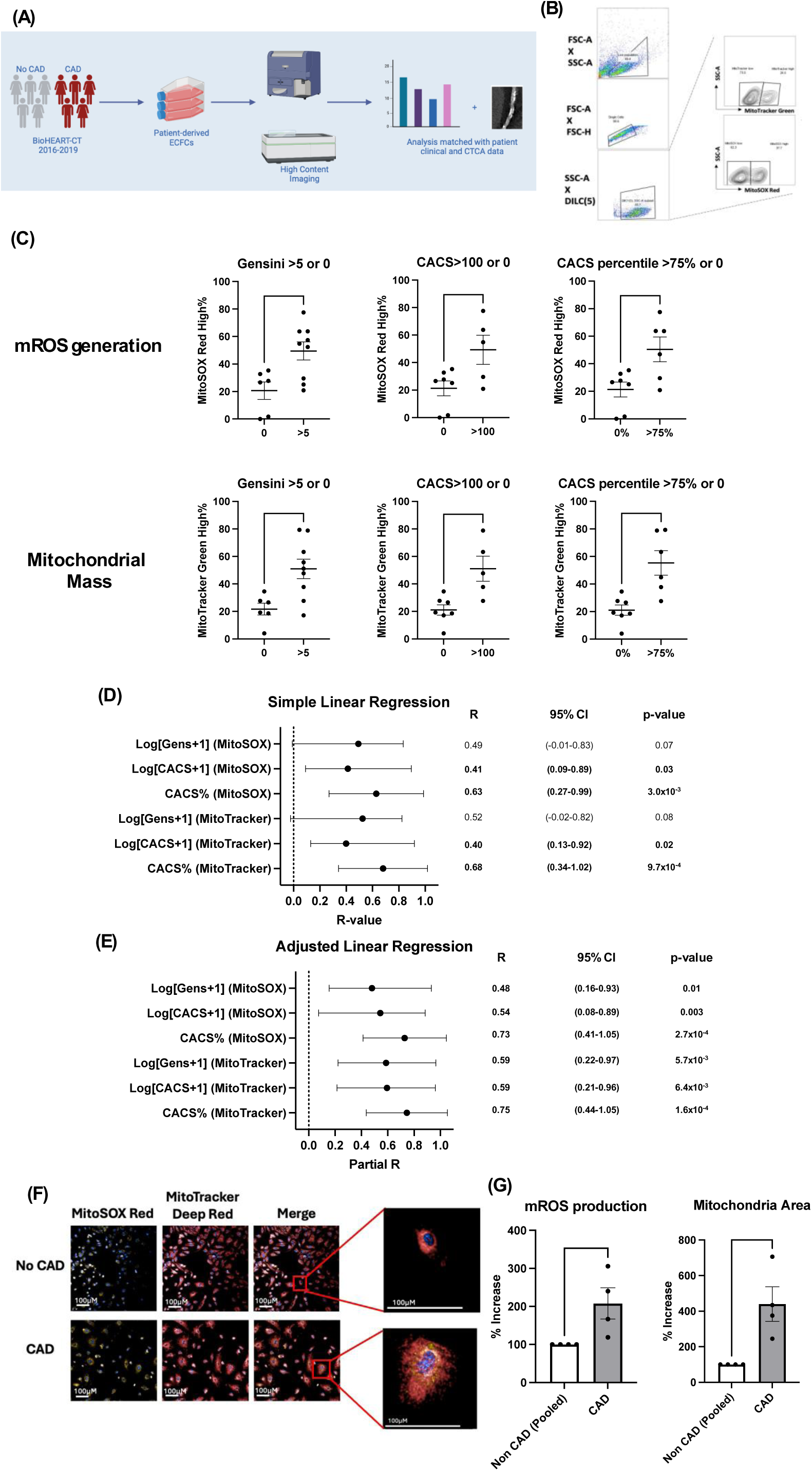
Mitochondrial function in ECFCs and its association with CAD Status. (A)Schematic diagram of mROS generation and mitochondrial mass characterisation experimental workflow.(B)Representative gating strategy for flow cytometry of MitoSOX Red and MitoTracker Green. MitoSOX Red and MitoTracker Green analyses were used to measure mROS production and mitochondrial mass. Results are expressed as either percentage of MitoSOX Red^High^ or MitoTracker Green^High^.(C)mROS production and mitochondrial mass between CAD (Gensini>5, CACS>100AU or >75% percentile) and non-CAD (Gensini=0, CACS=0AU or 0% percentile) patients. Results are presented as mean±SEM.(D-E)Forest plot of (D)simple and (E)adjusted (age, sex, diabetes, hypertension, hypercholestermia and smoking status) linear regression analyses performed for MitoSOX Red^High^ and MitoTracker Green^High^ proportions against CAD Severity (Log[Gensini+1], Log[CACS+1], CACS percentile). Error bars represent 95% CI.(F)Representative images of mitochondrial signatures from CAD and non-CAD ECFCs analysed by the Opera Phenix-method at 20x magnification(Scale bar=100μM).(G)mROS production and mitochondria area between CAD and non-CAD patients. Results are presented as mean±SEM. Statistical analyses were performed using non-parametric or parametric t-test or Firth’s bias-reduced penalized likelihood method for logistic regression test.*P<5.0×10^−2^;**P<1.0×10^−2^. CACS, coronary artery calcium score;mROS, mitochondrial reactive oxygen species.

### Multimodal analyses show strong associations between coronary calcium score/burden and ECFC mitochondrial dysregulation

Subsequently, we explored if a linear relationship existed between proportion of high mROS-generating and high mitochondrial-mass ECFCs with CAD severity. Both unadjusted and adjusted linear regressions were performed to adjust for covariates (age, sex and SMuRFs;Figure 3D-E). With simple linear regression, both MitoSOX Red^high^ (R=0.41, P=3×10^−2^) and MitoTracker Green^high^ (R=0.40, P=2×10^−2^) expressing ECFCs had strong correlations with Log[CACS+1]. When compared against CACS percentile, the strength of MitoSOX Red^high^ (R=0.63, P=3.0×10^−3^) and MitoTracker Green^high^ (R=0.68, P=9.7×10^−4^) increased significantly. Log[Gensini+1] did not show a significant correlation for proportionality of high mROS-generation or mitochondrial-mass. Subsequently, we adjusted for age, sex and presence of SMuRFs covariates (Figure 3E). Here, all metrics of CAD severity exhibited strong correlations when compared to MitoSOX Red^high^ and MitoTracker Green^high^ ECFCs (P<5.0×10^−2^). In particular, CACS percentile against MitoSOX Red^high^ (R=0.73, P=2.7×10^−4^) and MitoTracker Green^high^ (R=0.75, P=1.6×10^−4^) had very strong correlations. Logistic regression was also performed however no significant odds was elicited in both unadjusted and adjusted models (Supp. Fig 3A). Nevertheless, these findings support and build on our previous observation that patient-derived ECFCs maintain dyregulated mitochondrial redox state reflective of their CAD clinical phenotype. Next, we applied our recently developed Opera Phenix method^19^ to quantify and validate mitochondrial redox dysregulation in patient-derived ECFCs (Figure 3F). Consistent with the flow cytometry analyses, we observed that CAD-ECFCs exhibited a 108% higher mROS production (P=4.0×10^−2^), and 340% increased mitochondria area (P=1.0×10^−2^;Figure 3G).

### OxLDL is preferentially internalised by CAD ECFCS

Given the established role of oxidised LDL (oxLDL) in atherosclerosis and endothelial mitochondria dysfunction^19, 24^, we assessed the possibility of differential oxLDL-uptake between CAD and non-CAD ECFCs. Notably, CAD-ECFCs exhibited a 2.5-fold increase in oxLDL uptake compared to non-CAD ECFCs (P=2.9×10⁻²;Figure 4A). However, expression levels of key lipid transport receptors (*CD36, ABCA1* or *ABCG1*) were not significantly different between groups (Supp. Figure 3A-C), whilst *LOX-1* was not expressed at all in ECFCs (Supp. File 1)^25, 26^, suggesting alternate receptors/mechanisms of endothelial oxLDL-uptake.

**Figure 4.**
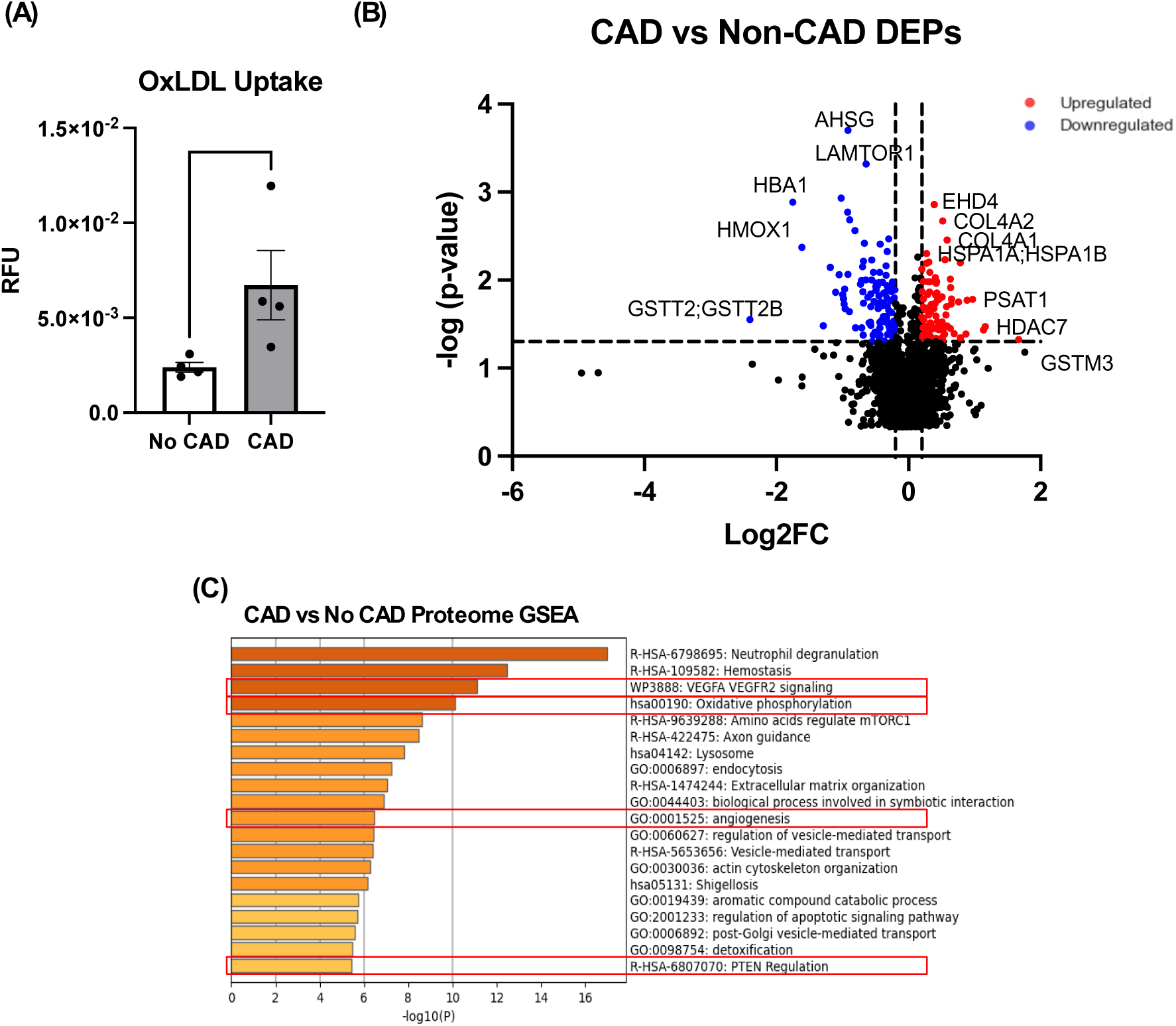
CAD ECFCs have significantly increased oxLDL uptake. (A)oxLDL uptake of CAD(n=4) or non-CAD ECFCs(n=4).Results are presented as mean±SEM.(B)DEPs between CAD(n=3) and non-CAD(n=3) ECFCs after exposure to oxLDL.(C)Enrichment analysis of DEPs from the proteome of CAD and non-CAD ECFCs.Bars=median.ns=no significance, *P<5.0×10^−2^.Statistical analyses were performed using Mann-Whitney U-test or independent samples t-test with FDR correction(FDR<0.05). OxLDL, oxidised low-density lipoprotein;DEP;differentially expressed proteins.

To investigate this further, we performed cellular proteomic profiling on oxLDL-treated ECFCs using micro-HPLC-qTOF-MS and DirectDIA+ with the BGS workflow (Figure 3B). A total of 194 differentially expressed proteins (DEPs) were identified between CAD and non-CAD groups, with 105 proteins significantly downregulated in non-CAD samples, representing 8.51% of the total proteome (Supp. File 2). Interestingly, there were no overlap between DEGs and DEPs. Again, no differences in influx and efflux receptors were found (Supp. File 2). GO enrichment anaylsis highlighted several key pathways including VEGFA/VEGFR2, oxidative phosphorylation, angiogenesis and PTEN regulatory pathways (Figure 4C;Supp. File 3), indicating distinct molecular adaptations in CAD ECFCs that may underlie enhanced oxLDL-uptake.

### Endothelial *CCBE1* Knockdown Does Not Alter Endothelial Function but Induces Mitochondrial Morphological Remodeling

Next, we explored the temporal expression of *CCBE1* with oxLDL-treated HUVECs (Figure 5A). We found *CCBE1* expression increased five-fold (P<5.0×10^−2^) at the 72h time point (Figure 5B). This occurred independently of canonical endothelial inflammatory markers, *VCAM-1* and *ICAM-1* expression (Supp. Figure 5A-B), suggesting that oxLDL may trigger a distinct metablic response in ECs. Silencing *CCBE1* in basal conditions was not feasible due to its minimal expression; however in oxLDL-stimulated HUVECs, *CCBE1* expression was successfully decreased by 89% at the mRNA level (P<1.0×10^−4^, Supp. Figure 5C-E), coinciding with the increase in *CCBE1* expression post-oxLDL exposure. When *CCBE1* was silenced, no impact on cell viability was observed (Supp. Figure 5F). Notably, *CCBE1* knockdown had no significant impact on the expression of lipid transporters (*CD36, ABCA1* and *ABCG1*) or inflammatory markers (*VCAM-1, ICAM-1*;Supp. Figure 5G-K). Protein-level suppression of CCBE1 was more modest, with only a 22% decrease observed (P=3.0×10^− 2^;Supp. Figure 5L).

**Figure 5.**
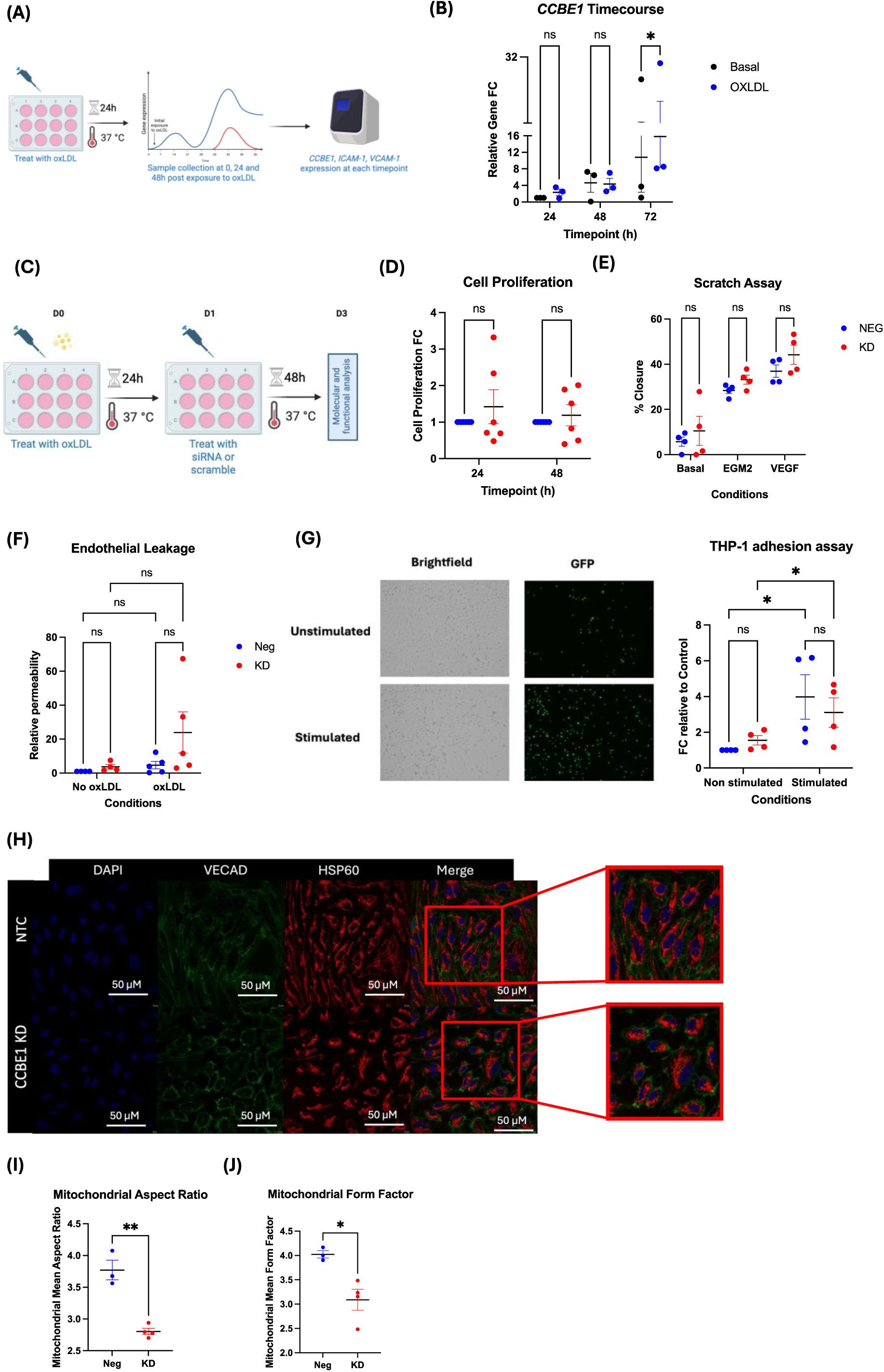
*CCBE1* knockdown does not affect endothelial cell function but induces significant morphological changes to the endothelial mitochondria. (A)Schematic diagram of experimental workflow for investigating *CCBE1, VCAM-1, ICAM-1* gene expression after oxLDL exposure.(B)Time course of CCBE1 expression by qRT-PCR in HUVECs treated with oxLDL(32.5 μg/ml).(C)Schematic diagram of experimental workflow for investigating *CCBE1* knockdown effect.(D)Cell proliferation and (E)Migration was following treatment with si*CCBE1* in oxLDL-treated HUVECs.(F)Endothelial leakage following treatment with si*CCBE1* in oxLDL-treated HUVECs.(G)Representative confocal images and analysis of THP-1 adhering to oxLDL-treated HUVECs with or without TNF-⍺ stimulation (H)Representative confocal images endothelial cell marker VE-Cadherin and mitochondrial marker HSP60(Scale bar=50μM). Image analysis results of (I)Mitochondrial aspect ratio, (J)Mitochondrial form factor.Paired samples t-test was performed for these experiments.Results are presented as mean±SEM.ns=no significance, *P<5.0×10^− 2^, **P<1.0×10^−2^. OxLDL, oxidised low-density lipoprotein.

To evaluate the functional relevance of CCBE1 in EC biology, we assessed its effect on cell proliferation, migration, barrier integrity and monocyte adhesion (Figure 5C). *CCBE1* knockdown had no impact on HUVEC proliferation, migration (Figure 5D-E;Supp. Figure 6A), or monolayer permeability (Figure 5F). There was no significant difference in THP-1 adhesion under both basal and TNF-⍺-stimulated conditions, consistent with the absence of changes in *VCAM-1* and *ICAM-1* expression following *CCBE1* knockdown (Figure 5G;Supp. Figure 6B). Collectively, these findings suggest that *CCBE1* expression is a downstream response to oxLDL exposure in ECs, potentially reflecting non-inflammatory metabolic reprogramming, but its knockdown does not affect classical endothelial functions or lipid handling under these conditions.

Subsequently, to elucidate a potential role for CCBE1 in mitochondrial function, we first confirmed its subcellular localisation in ECFCs. Western blot analysis of subcellular protein fractions revealed that CCBE1 is distributed across the mitochondrial, cytosolic, membrane, and nuclear compartments (Supp. Figure 6C). This broad subcellular distribution, including enrichment in the mitochondrial fraction, prompted us to investigate whether CCBE1 influences mitochondrial structure.

To this end, we examined the effects of *CCBE1* knockdown on mitochondrial morphology. *CCBE1* silencing resulted in a a substantial shift in mitochondrial morphology, transitioning from a characteristic elongated structure to a small, rounded phenotype (Figure 5H). Quantitative analysis of confocal microscopy images revealed that, although total mitochondrial area remained unchanged (Supp. Figure 6C), both aspect-ratio (AR) and form-factor (FF) were significantly reduced, indicative of mitochondrial fragmentation. Specifically, AR decreased by 26% (P=1.1×10⁻³;Figure 5I), whilst FF decreased by 23% (P=1.6×10⁻²;Figure 5J). These findings indicate a disruption in mitochondrial network integrity, although no significant changes were observed in mitochondrial branching (Supp. Figure 6D–H).

### Endothelial *CCBE1* Regulates Mitochondrial Metabolism and Bioenergetic Adaptations

Given the observed shift in mitochondrial morphology following *CCBE1* knockdown, we next investigated its impact on mitochondrial metabolism and bioenergetic function. To explore potential mechanistic contributors, we first examined the expression of genes associated with mitochondrial biogenesis, dynamics, and DNA maintenance. *CCBE1* knockdown significantly upregulated the expression of *TOMM20* (Translocase of Outer Mitochondrial Membrane 20), a marker of mitochondrial biogenesis (1.08-fold [95% CI:0.61–1.55];adjusted P=4.0×10⁻²;Figure 6A). However, no significant differences were observed in the expression of electron transport chain (ETC) components (*MTND1, SDHa, ATP5F1A, CYC1, COX6C*;Supp. Figure 7A) or mitochondrial DNA copy number (mtDNA-CN;Supp. Figure 7B).

**Figure 6.**
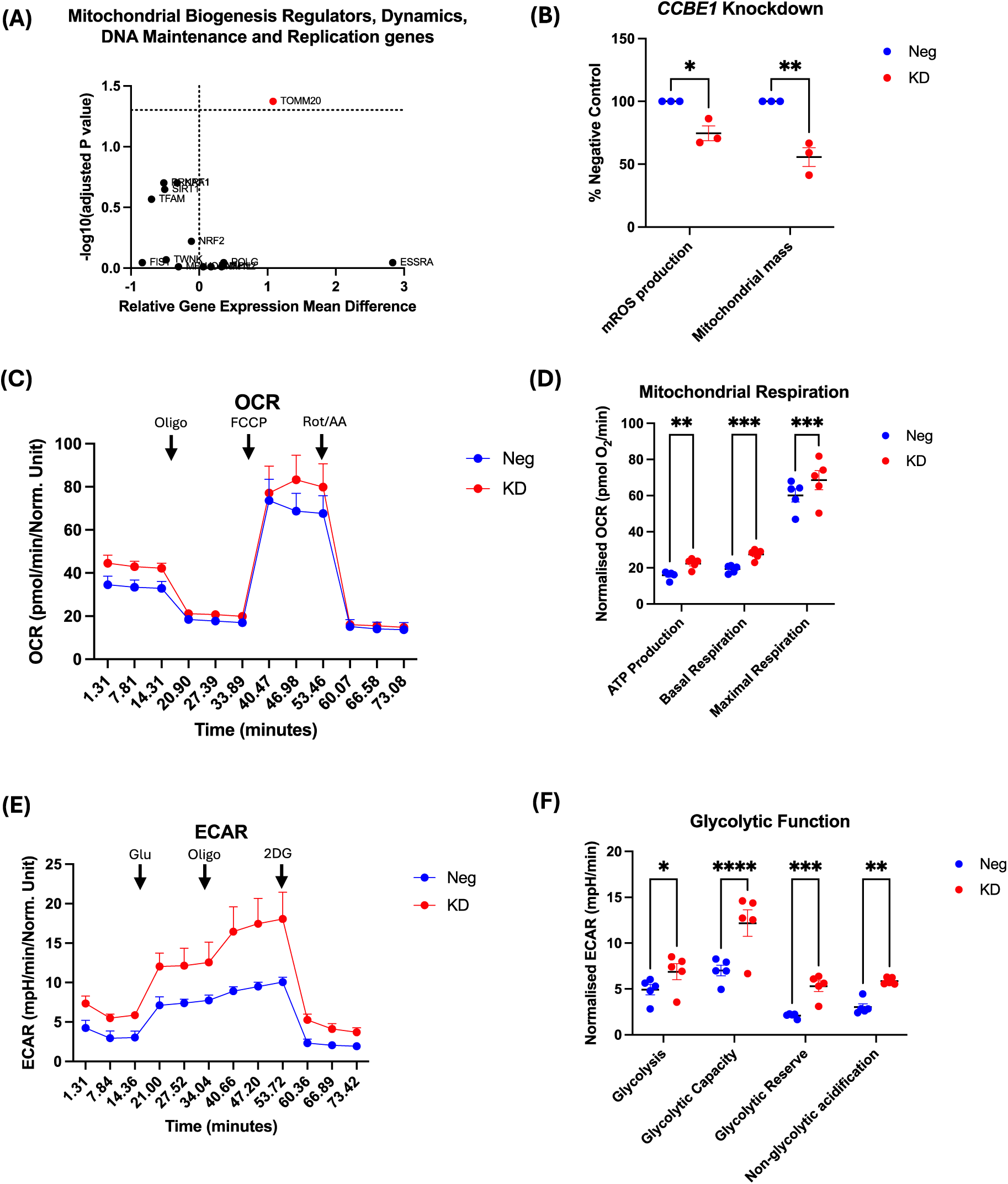
Endothelial *CCBE1* KD improves endothelial mitochondrial function. (A)Volcano Plot of genes associated with mitochondrial Biogenesis Regulators, mitochondrial Dynamics, mitochondrial DNA Maintenance and replication as determined by qRT-PCR(adjusted P-value<5.0×10^−2^).(B)mROS production and mitochondrial mass of oxLDL-treated HUVECs following Negatice control or si*CCBE1* treatment as determined by flow cytometry.(C)OCR curve of HUVECs treated with Negative control or si*CCBE1*.(D)ATP Production, basal respiration and maximal respiration were significantly increased after Negative control or si*CCBE1* treatment as determined by Seahorse analyser.(E)ECAR curve of HUVECs treated with Negative control or si*CCBE1*.(F)Glycolysis, Glycolytic capacity, Glycolytic reserve and Non-glycolytic acidification were significantly increased after Negative control or si*CCBE1* treatment as determined by Seahorse analyser.Results are presented as mean±SEM.Statistical analyses was performed used two-way ANOVA or multiple t-tests with FDR correction.FDR<0.05 was considered significant.*P<5.0×10^−2^, **P<1.0×10^−2^, ***P<1.0×10^−3^, ****P<1.0×10^−4^. OCR, oxygen consumption rate;ECAR, extracellular acidification rate.

Subsequently, we observed that *CCBE1* knockdown significantly reduced mROS production by 25% (P=3.9×10^−2^) and mitochondrial mass by 44% compared with the negative control (P=5.5×10^−3^;Figure 6B), indicating a more efficient and less stressed mitochondrial phenotype. Interestingly, in oxLDL-stimulated ECs–where *CCBE1* expression is upregulated–*CCBE1* silencing led to significant alterations in the mitochondrial profile (Figure 6C), including a 40% increase in ATP production (P=3.9×10⁻^3^), a 42% elevation in basal respiration (P=7.0×10^−4^), and a 14% increase in maximal respiration (P=5.0×10^− 4^;Figure 6D). These indicate a profound shift in endothelial bioenergetics and an adaptive response to mitochondrial remodelling. Despite these changes, proton leakage, spare respiratory capacity, and coupling efficiency remained unchanged (Supp. Figure 7C-E).

In addition to the reduction in mROS, increase in ATP production, and elevation in basal respiration with *CCBE1* knockdown, we also observed an increase in glycolytic ability (Figure 6E). ECs experienced significant mean increases in glycolysis (39.84%;P=3.9×10^−2^), glycolytic capacity (73.50%;P<1.0×10^−4^), glycolytic reserve (152.38%;P=8.70×10^−4^) and non-glycolytic acidification (93.73%;P=2.5×10^−3^) following *CCBE1* silencing (Figure 6F). Collectively, these data suggest that *CCBE1* knockdown promotes a metabolically active and adaptive endothelial phenotype characterised by enhanced bioenergetics and reduced mitochondrial oxidative stress.

### *CCBE1 cis*-eQTLs Study Reveals Strong Associations with CAD

Given prior evidence linking elevated *CCBE1* expression in CAD-ECFCs, we investigated associations between candidate *CCBE1 cis*-expression quantitative trait loci (*cis-*eQTLs) and CAD-related phenotypes. From the Stockholm-Tartu Atherosclerosis Reverse Network Engineering Task (STARNET) study^27^, we initially identified 589 candidate *cis*-eQTLs. This list was refined to 376 variants based on genotype availability and quality control in the BioHEART-CT cohort (n=976;Figure 7A).

**Figure 7.**
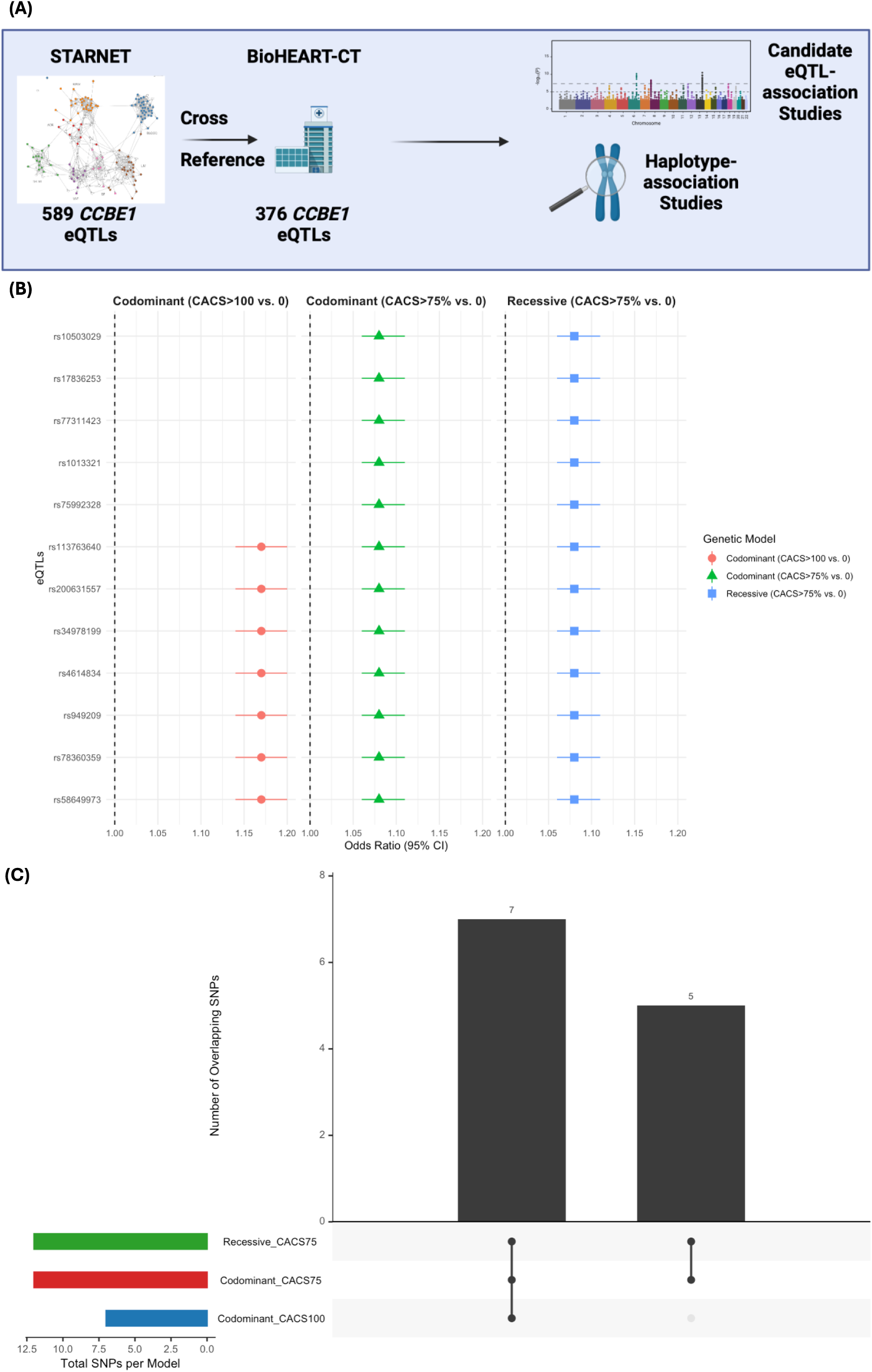
*CCBE1 cis-*eQTLs Strongly Predict CAD. (A)Schematic diagram of analysis workflow for investigating *CCBE1 cis*-eQTLS and haplotypes.(B)*cis*-eQTL-association analysis in a co-dominant and recessive adjusted-models of CACS (>100AU vs. 0AU) and CACS percentile (>75% vs. 0%).(C)UpSet plot showing shared *cis*-eQTLs between the different genetic models and disease metrics. Statistical analyses were performed using Firth’s bias-reduced penalized likelihood method for logistic regression test with FDR correction. FDR<0.05 was considered signficant. AU, Agatston unit;CACS, coronary artery calcium scoring;eQTL, expression quantitative trait loci.

We evaluated four genetic models—additive, dominant, recessive, and co-dominant—to assess associations between *CCBE1 cis*-eQTLs and CACS burden, using both CACS>100AU and >75th percentile as outcomes. No significant associations were observed in the additive or dominant models under either univariate or covariate-adjusted analyses (Supp. File 4). Similarly, the univariate co-dominant model showed no association. However, after adjustment for age, sex and SMuRF covariates, the co-dominant model revealed significant associations with >100AU. Individuals homozygous for the minor allele at rs58649973, rs78360359, rs949209, rs4614834, rs34978199, rs200631557 and rs113763640 showed increased odds of CACS>100 AU (OR=1.17;95% CI:1.14–1.20;adjusted P=6.2×10⁻²⁹;Figure 7B). All variants were intronic except rs4614834 (downstream). Notably, no association was detected in the recessive model, suggesting a genotype-specific effect unique to the co-dominant model.

Analysis of the >75% percentile outcome revealed a similar pattern. Additive and dominant models showed no significant associations in either univariate or adjusted analyses. However, covariate-adjusted models revealed significant associations in both the co-dominant and recessive frameworks. A shared set of *cis*-eQTLs— rs200631557, rs113763640, rs949209, rs75992328, rs1013321, rs77311423, rs58649973, rs78360359, rs17836253, rs10503029, rs34978199 and rs4614834—were associated with increased odds of CACS percentile>75% in Recessive model (OR=3.35;95% CI:2.21–5.08;adjusted P=3.4×10⁻⁷) and Co-dominant models (OR=3.30;95% CI:2.20– 5.10;adjusted P=4.0×10⁻⁷;Figure 7B). Except for rs34978199 (enhancer) and rs4614834 (downstream), all were intronic. Notably, all *cis-*eQTLs associated with CACS>100 AU were also linked to the >75th percentile outcome (Figure 7C).

Finally, we performed haplotype analysis based on linkage disequilibrium (LD) structure and defined haplotype blocks (Supp. Figure 8A-B). Bayesian logistic regression revealed a significant protective haplotype association with both CAD endpoints, consistent across univariate (Supp. Figure 7C-D;Supp. Table S8-9) and adjusted models (Supp. Figure 7E-F;Supp. Table S10-11).

## Discussion

Whilst endothelial signalling pathways has a well-recognised role in vascular health and pathogenesis of atherosclerosis, there is limited knowledge about what drives individual susceptibility or resilience to endothelial dysfunction and CAD. The BioHEART-CT study offers unique platform to investigate these mechanisms by combining two key strengths: firstly, a large-scale, deeply-phenotyped clinical cohort with advanced CT imaging to quantify coronary atherosclerosis burden; and secondly, a novel bank of patient ECFCs which we have previously shown to exhibit a phenotype reflective of their clinical CAD status^18^. This study supports our previous finding that patient-derived ECFCs retain a phenotype that reflects their CAD status^18, 19, 28^ even in *ex-vivo* conditions.

Transcriptomic analyses show significant dysregulation of mitochondrial-related genes, particularly those associated with ATP synthesis and the ETC (Figure 1C), consistent with our previous reports linking mitochondrial dysfunction to endothelial impairment^28^. However, it remains unclear whether these DEGs are causal factors in atherosclerosis or merely a consequence of the disease. Through an unbiased approach, we identified several DEGs present in ECFCs between CAD and non-CAD participants, with *CCBE1* emerging as a top candidate. Notably, a subset of non-CAD ECFCs exhibited high *RFLNA* expression, a regulator of filamin protein A (*FLNA*)—a gene implicated with atherosclerosis^29^—potentially indicating a protective role for *RFLNA* which should be investigated further. CCBE1 is known for to regulate lymphactic vessel development, acting as a mediator upstream of *VEGFC*^30, 31^. Its strong upregulation in CAD-ECFC’s in both study cohorts prompted us to prioritise CCBE1 for functional studies. Although CCBE1 has not been linked previously to CAD, its emerging roles in extracellular matrix remodelling^32^, suggest a novel causal link to endothelial dysfunction in atherosclerosis and CAD.

ECFCs contribute to vascular repair^33^, and *CCBE1* upregulation in CAD-ECFCs may reflect their activation to subclinical endothelial injury or signify intrinsic susceptibility to vascular dysfuction^34, 35^. Our findings from *cis*-eQTL analyses supports this idea that inherited variation in *CCBE1* locus/intronic variants is associated with increased coronary artery calcification and traditional CAD risk metrics (Figure 7B). Notably, this is the first report linking *CCBE1 cis*-eQTLs to CAD risk, supporting a role for CCBE1 as a biomarker of inherited endothelial vulnerability. This is important because of its selective expression in CAD-ECFCs making CCBE1 a promising diagnostic candidate for identifying individuals at increased CAD risk, particularly those without SMuRFs.

Consistent with prior studies, we observed a direct correlation between increased mitochondrial biogenesis and mROS levels in CAD-ECFCs (Figure 3)^36, 37^, which also exhibited increased oxLDL-uptake compared to non-CAD ECFCs (Figure 4A). This novel finding of enhanced oxLDL-uptake in CAD-ECFCs represents a potentially important biomarker of endothelial vulnerability and has not been previously reported. Mitochondrial dynamics are key regulators of mROS production; while mitochondrial fission has been linked to excess ROS production^38, 39^, excessive mitochondrial fusion—resulting in elongated, enlarged mitochondria—can similarly contribute to oxidative stress through impaired turnover and reduced metabolic adaptability^40^. These findings support a model in which enhanced oxLDL-uptake and dysregulated mitochondrial remodeling together drive endothelial oxidative dysfunction in CAD.

Our study provides mechanistic evidence that CCBE1 contributes to oxLDL-induced mitochondrial dysfunction in ECs, advancing our understanding into non-canonical pathways contributing to CAD susceptibility. In a physiologically-relevant model of oxLDL-stimulated ECs—a key mediator of vascular inflammation and CAD progression^41^-we demonstrate that oxLDL robustly induced *CCBE1* expression, independently of classical inflammatory adhesion molecules such as VCAM-1 and ICAM-1 (Figure 5B;Supp. Figure 5J–K)^42^. Functional experiments revealed that *CCBE1* knockdown conferred protection against oxLDL-induced dysfunction of mROS production and mitochondrial mass, whilst promoting a hyper-metabolic EC phenotype, with increased mitochondrial respiration and glycolysis (Figure 6B-E). This was accompanied by *TOMM20* upregulation, suggestive of enhanced protein import^43^ as a compensatory response to preserve mitochondrial mass which supports a role for CCBE1 in modulating mitochondrial function at a transcriptional level. Despite unchanged mitochondrial dynamic gene expression, *CCBE1* knockdown altered mitochondrial morphology, reducing aspect ratio and form factor (Figure 5H-I).

These functional and morphological findings diverge from previously reported CCBE1-TGFβ-DRP1 interactions in hepatocellular carcinoma^44^, suggesting a distinct, context-dependent role for CCBE1 in regulating mitochondrial structure and function in ECs. Collectively, our findings support a model in which CCBE1 acts downstream of oxLDL exposure to influence mitochondrial metabolism in ECs, independent of canonical inflammatory pathways^45^. Rather than being a primary driver of dysfunction, CCBE1 may represent an adaptive response to stress, facilitating endothelial metabolic remodelling under atherogenic conditions.

Our findings that *CCBE1* is upregulated in CAD-ECFCs compared to controls, along with functional studies supporting a causal role, have important relevance in the hunt for novel diagnostic and therapeutic targets. To evaluate *CCBE1*’s biomarker potential, future studies should evaluate its expression in circulating ECs and plasma-derived extracellular vesicles from CAD patients. Additionally, integrating *CCBE1* expression into multi-omic risk stratification models—including transcriptomic, proteomic, and metabolomics—may help clarify its potential added value beyond traditional CAD biomarkers. Longitudinal studies tracking *CCBE1* levels alongside coronary artery plaque progression will also be essential to establish its clinical utility in early detection and risk assessment.

Given *CCBE1’s* role in endothelial function and mitochondrial homeostasis, targeting this pathway may provide novel therapeutic strategies to enhance mitochondrial resilience in CAD-ECFCs, and offer endothelial protection. Furthermore, identifying small-molecule inhibitors or RNA-based therapeutics to modulate *CCBE1* expression could provide avenues for intervention. Given the association between *CCBE1* and oxLDL-induced stress, further investigation into lipid-lowering or antioxidant therapies in the context of *CCBE1* expression may also provide translational insights into CAD management.

Our study has several notable strengths. The use of CTCA to define CAD burden, as well as those with no angiographic evidence of atherosclerosis provided a much clearer clinical phenotype than available in general biobanks. This overcomes the challenges related to the substantial number of “healthy control” patients with subclinical atherosclerosis. The pairing of patient-derived ECFCs with this advanced imaging phenotype has facilitated our novel discovery, and supports the potential for larger scale use of this model for unravelling novel mechanisms of CAD susceptibility and resilience and for functional studies to validate potential causality.

Despite these strengths, our study has limitations.The pilot-scale human sample size limited statistical power and was underpowered for FDR-corrected analyses. Whilst functional knockdown experiments validated *CCBE1*’s role, future studies with large patient cohorts are needed to improve statistical robustness. Additionally, our functional studies were limited to *in-vitro* cell-based studies, necessitating validation *in-vivo*. Further insights could be gained with an EC-specific *CCBE1* knockout models, such as ApoE^−/-^ mice, potentially incorporating tandem stenosis of the carotid artery to better mimick human plaque progression^11, 46^. Finally, our transcriptomic results are specific to ECFCs, and CCBE1*’s* role in other EC subtypes remains to be explored.

In conclusion, the integration of *ex-vivo* ECFC transcriptomics profiling with CT-based plaque imaging has revealed novel mechansims contributing to atherosclerotic pathophysiology, offering tragets for prevention and treatment. Using a standardised approach to isolate ECFCs from peripheral blood, we demonstrated that *CCBE1* expression is elevated in CAD patients, particularly within the SMuRF-less cohort, and that its knockdown improved mitochondrial function in ECs. These findings suggest a previously unrecognised role for CCBE1 in CAD and atherosclerosis. Future studies are needed to determine where *CCBE1* modulation can be leveraged to prevent or treat atherosclerosis. Overall, the data from this study demonstrate the potential of ECFC-based transcriptomic profiling to facilitate new insights into the biology of coronary artery disease and identifying candidates for precision therapy.

## Acknowledgements

We acknowledge the participants, radiographers, nursing staff and Cardiovascular Discovery Group team members who have dedicated their time to the BioHEART study.

## References

1. Dawber TR, Moore FE, Mann GV. II. Coronary Heart Disease in the Framingham Study. Am J Public Health Nations Health. 1957;47:4–24.

2. Prediction of Coronary Heart Disease Using Risk Factor Categories | Circulation. Available at https://www.ahajournals.org/doi/10.1161/01.cir.97.18.1837?url_ver=Z39.88-2003&rfr_id=ori:rid:crossref.org&rfr_dat=cr_pub%20%200pubmed. Accessed August 26, 2022.

3. Piepoli MF, Hoes AW, Agewall S, Albus C, Brotons C, Catapano AL, Cooney M-T, Corrà U, Cosyns B, Deaton C, Graham I, Hall MS, Hobbs FDR, Løchen M-L, Löllgen H, Marques-Vidal P, Perk J, Prescott E, Redon J, Richter DJ, Sattar N, Smulders Y, Tiberi M, van der Worp HB, van Dis I, Verschuren WMM. 2016 European Guidelines on cardiovascular disease prevention in clinical practice. Eur Heart J. 2016;37:2315–2381.

4. Goff DC, Lloyd-Jones DM, Bennett G, Coady S, D’Agostino RB, Gibbons R, Greenland P, Lackland DT, Levy D, O’Donnell CJ, Robinson JG, Schwartz JS, Shero ST, Smith SC, Sorlie P, Stone NJ, Wilson PWF. 2013 ACC/AHA Guideline on the Assessment of Cardiovascular Risk. Circulation. 2014;129:S49–S73.

5. Kong G, Chin YH, Chong B, Goh RSJ, Lim OZH, Ng CH, Muthiah M, Foo R, Vernon ST, Loh PH, Chan MY, Chew NWS, Figtree GA. Higher mortality in acute coronary syndrome patients without standard modifiable risk factors: Results from a global meta-analysis of 1, 285, 722 patients. International Journal of Cardiology. 2023;371:432–440.

6. Figtree GA, Vernon ST, Hadziosmanovic N, Sundström J, Alfredsson J, Arnott C, Delatour V, Leósdóttir M, Hagström E. Mortality in STEMI patients without standard modifiable risk factors: a sex-disaggregated analysis of SWEDEHEART registry data. The Lancet. 2021;397:1085–1094.

7. Vernon ST, Coffey S, D’Souza M, Chow CK, Kilian J, Hyun K, Shaw JA, Adams M, Roberts-Thomson P, Brieger D, Figtree GA. ST-Segment–Elevation Myocardial Infarction (STEMI) Patients Without Standard Modifiable Cardiovascular Risk Factors—How Common Are They, and What Are Their Outcomes? J Am Heart Assoc. 2019;8:e013296.

8. Vernon ST, Coffey S, Bhindi R, Soo Hoo SY, Nelson GI, Ward MR, Hansen PS, Asrress KN, Chow CK, Celermajer DS, O’Sullivan JF, Figtree GA. Increasing proportion of ST elevation myocardial infarction patients with coronary atherosclerosis poorly explained by standard modifiable risk factors. European Journal of Preventive Cardiology. 2017;24:1824–1830.

9. Figtree GA, Vernon ST. Coronary artery disease patients without standard modifiable risk factors (SMuRFs)- a forgotten group calling out for new discoveries. Cardiovascular Research. 2021;117:e76–e78.

10. Steven S, Frenis K, Oelze M, Kalinovic S, Kuntic M, Bayo Jimenez MT, Vujacic-Mirski K, Helmstädter J, Kröller-Schön S, Münzel T, Daiber A. Vascular Inflammation and Oxidative Stress: Major Triggers for Cardiovascular Disease. Oxidative Medicine and Cellular Longevity. 2019;2019:7092151.

11. Lee WE, Genetzakis E, Figtree GA. Novel Strategies in the Early Detection and Treatment of Endothelial Cell-Specific Mitochondrial Dysfunction in Coronary Artery Disease. Antioxidants. 2023;12:1359.

12. Boulanger CM. Endothelium. Arteriosclerosis, Thrombosis, and Vascular Biology. 2016;36:e26–e31.

13. Esper RJ, Nordaby RA, Vilariño JO, Paragano A, Cacharrón JL, Machado RA. Endothelial dysfunction: a comprehensive appraisal. Cardiovascular Diabetology. 2006;5:4.

14. Mudau M, Genis A, Lochner A, Strijdom H. Endothelial dysfunction: the early predictor of atherosclerosis. Cardiovasc J Afr. 2012;23:222–231.

15. Barthelmes J, Nägele MP, Ludovici V, Ruschitzka F, Sudano I, Flammer AJ. Endothelial dysfunction in cardiovascular disease and Flammer syndrome—similarities and differences. EPMA Journal. 2017;8:99–109.

16. Widder JD, Fraccarollo D, Galuppo P, Hansen JM, Jones DP, Ertl G, Bauersachs J. Attenuation of Angiotensin II-induced Vascular Dysfunction and Hypertension by Overexpression of Thioredoxin-2. Hypertension. 2009;54:338–344.

17. Kott KA, Vernon ST, Hansen T, Yu C, Bubb KJ, Coffey S, Sullivan D, Yang J, O’Sullivan J, Chow C, Patel S, Chong J, Celermajer DS, Kritharides L, Grieve SM, Figtree GA. Biobanking for discovery of novel cardiovascular biomarkers using imaging-quantified disease burden: protocol for the longitudinal, prospective, BioHEART-CT cohort study. BMJ Open. 2019;9:e028649.

18. Besnier M, Finemore M, Yu C, Kott KA, Vernon ST, Seebacher NA, Genetzakis E, Furman A, Tang O, Davis RL, Hansen T, Psaltis PJ, Bubb KJ, Wise SG, Grieve SM, Di Bartolo BA, Figtree GA. Patient Endothelial Colony-Forming Cells to Model Coronary Artery Disease Susceptibility and Unravel the Role of Dysregulated Mitochondrial Redox Signalling. Antioxidants. 2021;10:1547.

19. Lee WE, Besnier M, Genetzakis E, Tang O, Kott KA, Vernon ST, Gray MP, Grieve SM, Kassiou M, Figtree GA. High-Throughput Measure of Mitochondrial Superoxide Levels as a Marker of Coronary Artery Disease to Accelerate Drug Translation in Patient-Derived Endothelial Cells Using Opera Phenix® Technology. International Journal of Molecular Sciences. 2024;25:22.

20. Agatston AS, Janowitz WR, Hildner FJ, Zusmer NR, Viamonte M, Detrano R. Quantification of coronary artery calcium using ultrafast computed tomography. Journal of the American College of Cardiology. 1990;15:827–832.

21. Arnett DK, Blumenthal RS, Albert MA, Buroker AB, Goldberger ZD, Hahn EJ, Himmelfarb CD, Khera A, Lloyd-Jones D, McEvoy JW, Michos ED, Miedema MD, Muñoz D, Smith SC, Virani SS, Williams KA, Yeboah J, Ziaeian B. 2019 ACC/AHA Guideline on the Primary Prevention of Cardiovascular Disease: A Report of the American College of Cardiology/American Heart Association Task Force on Clinical Practice Guidelines. Circulation. 2019;140:e596–e646.

22. Hamilton-Craig CR, Chow CK, Younger JF, Jelinek VM, Chan J, Liew GY. Cardiac Society of Australia and New Zealand position statement executive summary: coronary artery calcium scoring. Medical Journal of Australia. 2017;207:357–361.

23. Murphy MP, Bayir H, Belousov V, Chang CJ, Davies KJA, Davies MJ, Dick TP, Finkel T, Forman HJ, Janssen-Heininger Y, Gems D, Kagan VE, Kalyanaraman B, Larsson N-G, Milne GL, Nyström T, Poulsen HE, Radi R, Van Remmen H, Schumacker PT, Thornalley PJ, Toyokuni S, Winterbourn CC, Yin H, Halliwell B. Guidelines for measuring reactive oxygen species and oxidative damage in cells and in vivo. Nat Metab. 2022;4:651–662.

24. Pirillo A, Norata GD, Catapano AL. LOX-1, OxLDL, and Atherosclerosis. Mediators of Inflammation. 2013;2013:152786.

25. Li D, Mehta JL. Intracellular Signaling of LOX-1 in Endothelial Cell Apoptosis. Circulation Research. 2009;104:566–568.

26. Mehta JL, Chen J, Hermonat PL, Romeo F, Novelli G. Lectin-like, oxidized low-density lipoprotein receptor-1 (LOX-1): A critical player in the development of atherosclerosis and related disorders. Cardiovascular Research. 2006;69:36–45.

27. Koplev S, Seldin M, Sukhavasi K, Ermel R, Pang S, Zeng L, Bankier S, Di Narzo A, Cheng H, Meda V, Ma A, Talukdar H, Cohain A, Amadori L, Argmann C, Houten SM, Franzén O, Mocci G, Meelu OA, Ishikawa K, Whatling C, Jain A, Jain RK, Gan L-M, Giannarelli C, Roussos P, Hao K, Schunkert H, Michoel T, Ruusalepp A, Schadt EE, Kovacic JC, Lusis AJ, Björkegren JLM. A mechanistic framework for cardiometabolic and coronary artery diseases. Nat Cardiovasc Res. 2022;1:85–100.

28. Lee WE, Genetzakis E, Barsha G, Vescovi J, Mifsud C, Vernon ST, Nguyen TV, Gray MP, Grieve SM, Figtree GA. Expression of Myeloperoxidase in Patient-Derived Endothelial Colony-Forming Cells—Associations with Coronary Artery Disease and Mitochondrial Function. Biomolecules. 2024;14:1308.

29. Bandaru S, Ala C, Salimi R, Akula MK, Ekstrand M, Devarakonda S, Karlsson J, Van den Eynden J, Bergström G, Larsson E, Levin M, Borén J, Bergo MO, Akyürek LM. Targeting Filamin A Reduces Macrophage Activity and Atherosclerosis. Circulation. 2019;140:67–79.

30. Bos FL, Caunt M, Peterson-Maduro J, Planas-Paz L, Kowalski J, Karpanen T, van Impel A, Tong R, Ernst JA, Korving J, van Es JH, Lammert E, Duckers HJ, Schulte-Merker S. CCBE1 Is Essential for Mammalian Lymphatic Vascular Development and Enhances the Lymphangiogenic Effect of Vascular Endothelial Growth Factor-C In Vivo. Circulation Research. 2011;109:486–491.

31. Jeltsch M, Jha SK, Tvorogov D, Anisimov A, Leppänen V-M, Holopainen T, Kivelä R, Ortega S, Kärpanen T, Alitalo K. CCBE1 Enhances Lymphangiogenesis via A Disintegrin and Metalloprotease With Thrombospondin Motifs-3–Mediated Vascular Endothelial Growth Factor-C Activation. Circulation. 2014;129:1962–1971.

32. Yuan Z, Li Y, Zhang S, Wang X, Dou H, Yu X, Zhang Z, Yang S, Xiao M. Extracellular matrix remodeling in tumor progression and immune escape: from mechanisms to treatments. Molecular Cancer. 2023;22:48.

33. Banno K, Yoder MC. Tissue regeneration using endothelial colony-forming cells: promising cells for vascular repair. Pediatr Res. 2018;83:283–290.

34. Xu Q. Progenitor cells in vascular repair. Current Opinion in Lipidology. 2007;18:534.

35. Xu Q. Mouse Models of Arteriosclerosis: From Arterial Injuries to Vascular Grafts. The American Journal of Pathology. 2004;165:1–10.

36. Zhang B, Pan C, Feng C, Yan C, Yu Y, Chen Z, Guo C, Wang X. Role of mitochondrial reactive oxygen species in homeostasis regulation. Redox Rep.;27:45–52.

37. Jang K-J, Mano H, Aoki K, Hayashi T, Muto A, Nambu Y, Takahashi K, Itoh K, Taketani S, Nutt SL, Igarashi K, Shimizu A, Sugai M. Mitochondrial function provides instructive signals for activation-induced B-cell fates. Nat Commun. 2015;6:6750.

38. Huang Q, Zhan L, Cao H, Li J, Lyu Y, Guo X, Zhang J, Ji L, Ren T, An J, Liu B, Nie Y, Xing J. Increased mitochondrial fission promotes autophagy and hepatocellular carcinoma cell survival through the ROS-modulated coordinated regulation of the NFKB and TP53 pathways. Autophagy. 2016;12:999–1014.

39. Sánchez-Alvarez R, De Francesco EM, Fiorillo M, Sotgia F, Lisanti MP. Mitochondrial Fission Factor (MFF) Inhibits Mitochondrial Metabolism and Reduces Breast Cancer Stem Cell (CSC) Activity. Front Oncol. 2020;10. doi:10.3389/fonc.2020.01776.

40. Yoon Y-S, Yoon D-S, Lim IK, Yoon S-H, Chung H-Y, Rojo M, Malka F, Jou M-J, Martinou J-C, Yoon G. Formation of elongated giant mitochondria in DFO-induced cellular senescence: Involvement of enhanced fusion process through modulation of Fis1. Journal of Cellular Physiology. 2006;209:468–480.

41. Hong CG, Florida E, Li H, Parel PM, Mehta NN, Sorokin AV. Oxidized low-density lipoprotein associates with cardiovascular disease by a vicious cycle of atherosclerosis and inflammation: A systematic review and meta-analysis. Front Cardiovasc Med. 2023;9:1023651.

42. Singh V, Kaur R, Kumari P, Pasricha C, Singh R. ICAM-1 and VCAM-1: Gatekeepers in various inflammatory and cardiovascular disorders. Clinica Chimica Acta. 2023;548:117487.

43. Su J, Tian X, Wang Z, Yang J, Sun S, Sui S-F. Structure of the intact Tom20 receptor in the human translocase of the outer membrane complex. PNAS Nexus. 2024;3:pgae269.

44. Tian G-A, Xu W-T, Zhang X-L, Zhou Y-Q, Sun Y, Hu L-P, Jiang S-H, Nie H-Z, Zhang Z-G, Zhu L, Li J, Yang X-M, Yao L-L. CCBE1 promotes mitochondrial fusion by inhibiting the TGFβ-DRP1 axis to prevent the progression of hepatocellular carcinoma. Matrix Biology. 2023;117:31–45.

45. Jiang J, Hiron TK, Chalisey A, Malhotra Y, Agbaedeng T, O’Callaghan CA. Ox-LDL induces a non-inflammatory response enriched for coronary artery disease risk in human endothelial cells. 2024;:2024.11.19.624299.

46. Chuaiphichai S, Chu SM, Carnicer R, Kelly M, Bendall JK, Simon JN, Douglas G, Crabtree MJ, Casadei B, Channon KM. Endothelial cell-specific roles for tetrahydrobiopterin in myocardial function, cardiac hypertrophy, and response to myocardial ischemia-reperfusion injury. American Journal of Physiology-Heart and Circulatory Physiology. 2023;324:H430–H442.

